# A model of ethanol self-administration in head-fixed mice

**DOI:** 10.1101/2024.02.17.580838

**Authors:** Amy L. Ward, Kion T. Winston, Sophie Buchmaier, Cynara J. Cooper, Rachel E. Clarke, Michael R. Martino, Kelsey M. Vollmer, Jacqueline E. Paniccia, Marcus S. Bell, Elizabeth M. Doncheck, Roger I. Grant, Joshua Boquiren, Jade Baek, Logan M. Manusky, Annaka M. Westphal, Lisa M. Green, Bayleigh E. Pagoota, James M. Otis, Jennifer A. Rinker

## Abstract

**Background:** Significant advances in neurotechnology, such as the application of two-photon (2P) imaging of biosensors *in vivo*, have enabled unparalleled longitudinal and high-resolution access to neural circuits that coordinate behavior in rodents. Integration of these techniques would be groundbreaking for the study of alcohol use disorder (AUD). AUD is rooted in significant neural adaptations that could be functionally monitored and manipulated at the single-cell level across development of dependence in rodents. However, 2P imaging and related methodologies often require or are facilitated by head-fixation, and a lack of head-fixed models have hindered their integration for the study of alcohol dependence.

**Methods:** We developed a head-fixed model in which animals learned to self-administer ethanol across ∼14 days. Active lever responding resulted in a tone cue and ethanol reward, whereas responding on the inactive lever resulted in neither cue nor ethanol reward. Following acquisition, animals extinguished lever pressing across a minimum of 10 days. And finally, animals were tested separately for both cue- and ethanol-induced reinstatement of lever pressing.

**Results:** Here we show, for the first time, that in our head-fixed ethanol self-administration model male and female mice reliably pressed an active, but not inactive, lever for an oral ethanol reward. Ethanol rewards positively correlated with blood ethanol concentrations, at pharmacologically relevant levels. Furthermore, mice extinguished ethanol self-administration when the ethanol reward and cue were omitted, suggesting active lever pressing was ethanol directed. Following extinction, presentation of the ethanol-associated cue or priming with ethanol itself invigorated reinstatement of ethanol seeking, modeling relapse in a manner that replicates decades of work in freely-moving rodent studies.

**Conclusions:** Overall, our head-fixed ethanol self-administration model will allow for incorporation of novel technologies that require or are greatly facilitated by head-fixation, improving our ability to study and understand the neural adaptations and computations that underlie alcohol dependence.

## INTRODUCTION

Alcohol use disorder (AUD) is characterized by continuous cycles of positive reinforcement, wherein the pleasant, euphoric effects of alcohol increase alcohol consumption (Brown et al., 1980) and negative reinforcement, wherein individuals consume alcohol to avoid withdrawal or alleviate anxiety (Cooper et al., 1995, Koob and Volkow, 2010). AUD is a chronic, relapsing neuropsychological condition where excessive alcohol seeking, and the inability to stop seeking despite negative consequences, are hallmarks of dependence (NIAAA, 2023). Repeatedly experiencing the positive reinforcing effects of alcohol can increase the likelihood of alcohol-seeking behaviors (Cho et al., 2019), and contextual associations, such as drug-associated cues, environments or the drug itself, may trigger relapse in individuals with AUD. To adequately replicate this reinforcement of seeking behaviors in preclinical models, researchers often use operant self-administration paradigms, which reinforce alcohol (ethanol) seeking (e.g., active lever presses or nose-pokes), the rewarding aspects of ethanol, and perpetuate a cycle that strengthens ethanol-seeking behavior in rodent models (Gass and Olive, 2007, June and Gilpin, 2010, Samson and Czachowski, 2003). During self-administration, animals learn an operant task which is subsequently reinforced by drug delivery, often paired with a discrete drug-associated cue, like a tone or a light (Haney and Spealman, 2008). Following learning, this reinforced behavior is extinguished through drug and cue omission. Once this behavior is adequately extinguished, animals can be tested for reinstatement of seeking behavior to a variety of stimuli, including cues and the drug itself, modeling relapse in humans (Epstein et al., 2006). Therefore, preclinical models can effectively parallel the complex cycle of substance use disorders through self-administration (Koob and Le Moal, 1997).

Considering the effectiveness of using operant conditioning as a behavioral model, pairing this approach with state-of-the-art techniques that enable improved interrogation of neural circuits would improve our ability to define the neurobiological mechanisms of AUD. For example, with the advent of biosensor technologies, two-photon (2P) microscopy could be applied to self-administration experiments, enabling longitudinal visualization of activity (e.g., through calcium or voltage sensors) and morphology (e.g., through synaptic fluorophores) in single cell-type specific neurons, dendrites, and axons (Chen et al., 2013, Grienberger et al., 2022, Svoboda and Yasuda, 2006). Until recently, 2P imaging was constrained by the relatively limited behavioral procedures amenable to head-fixation in mice, but creative adaptations have expanded the number of relevant behavioral paradigms. Specifically, recent developments have incorporated elements of operant conditioning (e.g., levers) and delivery mechanisms (e.g., sipper tubes for fluid delivery or IV catheterization for drug delivery) with sophisticated 2P microscopy, allowing for visualization of activity in single neurons from the onset of self-administration experiments through reinstatement (Vollmer et al., 2021, Vollmer et al., 2022, Paniccia et al., 2024). While recent advances have been made, enabling some aspects of ethanol-related behaviors to be studied in head-fixed animals (Kalelkar et al., 2024, Timme et al., 2024), a head-fixed model of operant ethanol self-administration has not yet been created or experimentally validated.

Here we present a model of ethanol self-administration in head-fixed mice, and we base our model design on decades of research using self-administration in freely moving mice (Meisch, 2001, Lopez and Becker, 2014). Within this paradigm, we find that head-fixed mice learn to press a rewarded active, but not unrewarded inactive, lever resulting in a tone cue followed by an ethanol reward. Mice rapidly discriminate between the levers and achieve physiologically relevant blood ethanol concentrations that are predicted by the number of ethanol deliveries. Moreover, when the cue and ethanol rewards are omitted during extinction, mice learn to suppress active lever pressing across days. Notably, when we re-introduce cue or ethanol rewards following extinction learning, we find that mice will significantly reinstate drug-seeking behaviors. Collectively, these results suggest that we can effectively model ethanol self-administration in head-fixed mice, establishing an effective paradigm for use with high resolution 2P microscopy to provide extensive insight into the neural mechanisms underlying AUD.

## MATERIALS AND METHODS

### Animals

Male and female mice from various strains [i.e., C57BL/6J (Jax Labs #000664), PV-2A-Cre (Jax Labs #012358), or wildtype C57BL/6J background mice from our breeding colony] at least 8 weeks old/20g minimum at study onset. Mice were group housed pre-operatively and single-housed postoperatively under a reversed 12:12-hour light cycle (lights off at 8:00AM; all experiments were performed during the dark phase) with access to standard rodent chow (Teklad Diet 2918, Inotiv) and water *ad libitum* except during ethanol self-administration sessions. Not all animals were exposed to every phase of experimentation, as some animals were utilized for other experiments but data was included for unmanipulated phases, thus the total sample size may vary. All animal experimental procedures were reviewed and approved by the Institutional Animal Care and Use Committee (IACUC) at the Medical University of South Carolina and adhere to NIH guidelines for working with laboratory animals (National Research Council, 2011).

### Head-Fixed Surgeries

Mice were anesthetized with isoflurane (0.8-1.5% in oxygen; 1L/min) and placed within a stereotaxic frame (Kopf Instruments) for intracranial surgeries. Ophthalmic ointment (Akorn), topical anesthetic (2% Lidocaine; Akorn), analgesic (Ketorlac, 2mg/kg, IP), and subcutaneous sterile saline (0.9% NaCl in water) treatments were given pre-operatively and intra-operatively for the health and pain management of the animal. Some animals received head-ring implantation only (see below) and utilized for behavior only (n=13), while some animals reported herein were utilized for other experimentation beyond the scope of this report and experienced additional surgical procedures (i.e. bilateral virus injection or fiber implantation for manipulations that occurred after the data reported were collected; n=25). A custom-made head-ring (stainless steel; 5 mm ID, 11 mm OD; Clemson Engineering) was implanted onto the animal skull to allow for head-fixation. Following surgeries animals received antibiotics (Cefazolin, 200mg/kg, sc) and recovered with access to food and water *ad libitum* for 14 days minimum.

### Ethanol Drinking Habituation

Following three to four weeks of recovery from surgery, animals were acclimated to ethanol (EtOH) consumption using a modified Drinking-in-the-Dark (DID) paradigm (Rhodes et al., 2005, Rhodes et al., 2007). Three hours into the dark cycle (lights off at 8AM), water bottles were replaced with volumetric sipper tubes containing 15% EtOH (v/v in reverse osmosis water, Decon Labs). Animals were permitted to drink for two hours each day over the course of seven days. Animals were weighed each day of home cage drinking and EtOH consumption was normalized to body weight (grams of EtOH consumed per kilogram of body weight; g/kg).

### Head-Fixed Ethanol Operant Self-Administration

#### Acquisition

To reduce possible stress caused by head-fixation, following home cage drinking mice underwent two days of habituation during which they were head-restrained in custom-built operant chambers for 45 minutes with no access to the levers, tone cue, or ethanol reward (for more details see (Vollmer et al., 2021). Once habituated to head-fixation, mice were trained on a fixed-ratio 1 (FR1) schedule of reinforcement operant conditioning paradigm. Behavioral systems interfaced with an Arduino and custom software developed in Python or MatLab (as described in (Vollmer et al., 2021)) were equipped to deliver ethanol reinforcers, record lever presses and collect lick contacts with either the liquid or the spout. During behavioral acquisition, the levers were placed in front of the mice. Pressing the active lever resulted in an immediate cue presentation (8kHz, 1.6s), and finally an ethanol reward. Mice self-administering ethanol received 15% EtOH droplets (∼3-5 µL) in front of their mouth. Reinforced active lever pressing resulted in a 20 second timeout period wherein continuous pressing of the active lever was recorded, but no reward or cue was presented. Pressing of the inactive lever presented neither cue nor reward. Mice underwent ethanol self-administration for 1-hour sessions where mice were allowed to freely self-administer ethanol for the entirety of the session.

#### Extinction & Reinstatement Testing

Following 14 days of self-administration, mice learned to extinguish prior ethanol-seeking behavior across for at least 10 days. During extinction, lever pressing no longer results in a cue or ethanol reward delivery. Following a minimum of 10 days of extinction, if animals have reached extinction criteria, which is defined as <20% of the average lever presses on the last two days of behavioral acquisition (Vollmer et al., 2021), they underwent either drug or cue reinstatement. For cue reinstatement, animals received a tone cue (8kHz, 2s) upon pressing the active lever but did not receive an ethanol reward. For drug reinstatement, animals were given access to a volumetric sipper tube filled with 15% ethanol for 30 minutes in their home cage prior to the reinstatement session. Between reinstatement sessions, mice underwent re-extinction (<20% of the average lever presses on the last two days of behavioral acquisition) to reestablish low lever pressing rates.

### Blood Ethanol Concentration (BEC) Measurements

To determine the degree of intoxication following operant self-administration behavioral sessions, retro-orbital blood samples were collected in heparinized capillary tubes in a subset of animals to measure blood ethanol concentrations (BECs). Blood samples were centrifuged at 10,000x*g* at 4°C for 10 minutes to separate red blood cells and plasma, and plasma samples were extracted and BEC determined using an Analox AM1 Analyzer (Analox Instruments).

### Data Collection and Statistical Analyses

The majority of pressing data during acquisition was collected using a custom MATLAB interface connected to an Arduino (Vollmer et al., 2021). A subset of pressing data (n=8) during acquisition was collected using a new, custom-made behavioral interface controlled by Python, also connected via an Arduino. Moreover, all licking behavior during acquisition was collected using the Python interface. GraphPad PRISM (Dotmatics, version 10) was used to conduct statistical analyses and, where appropriate, either analysis of variance (ANOVA; two-way or three-way) or paired t-tests were used to analyze behavioral data. The following independent variables were analyzed: lever presses (active or inactive), day (acquisition, extinction, or reinstatement), sex (male or female) and lick rate (early acquisition, late acquisition, active or timeout lever press).

## RESULTS

### Home Cage Ethanol Consumption

All mice were habituated to ethanol using a modified Drinking-in-the-Dark (DID) paradigm. Animals were given access to 15% EtOH (v/v) in their home cage via volumetric sipper tubes for 7 days a week prior to head-fixation (**Fig 2A**). Despite previous reports of sex-dependent differences in ethanol consumption (Rivera-Irizarry et al., 2023), we found that male and female mice drank similar amounts during the home cage access period [female mean 1.758 g/kg ± 0.18 vs male mean 1.98 g/kg ± 0.15 (**Fig 2A**); main effect of sex, F_1,36_=0.88, *p*=0.3526]. There was a main effect of day, F_3.405,122.0_=4.135, *p*=0.0057, and Tukey’s post-hoc analyses showed that drinking on day 4 decreased compared to day 2 (*p*=0.0364), but stabilized after that, and there was no sex x day interaction, F_6,215_=1.280, *p*=0.2675. Please note, one drinking value during home-cage drinking was removed because of spillage.

**Figure 1.**
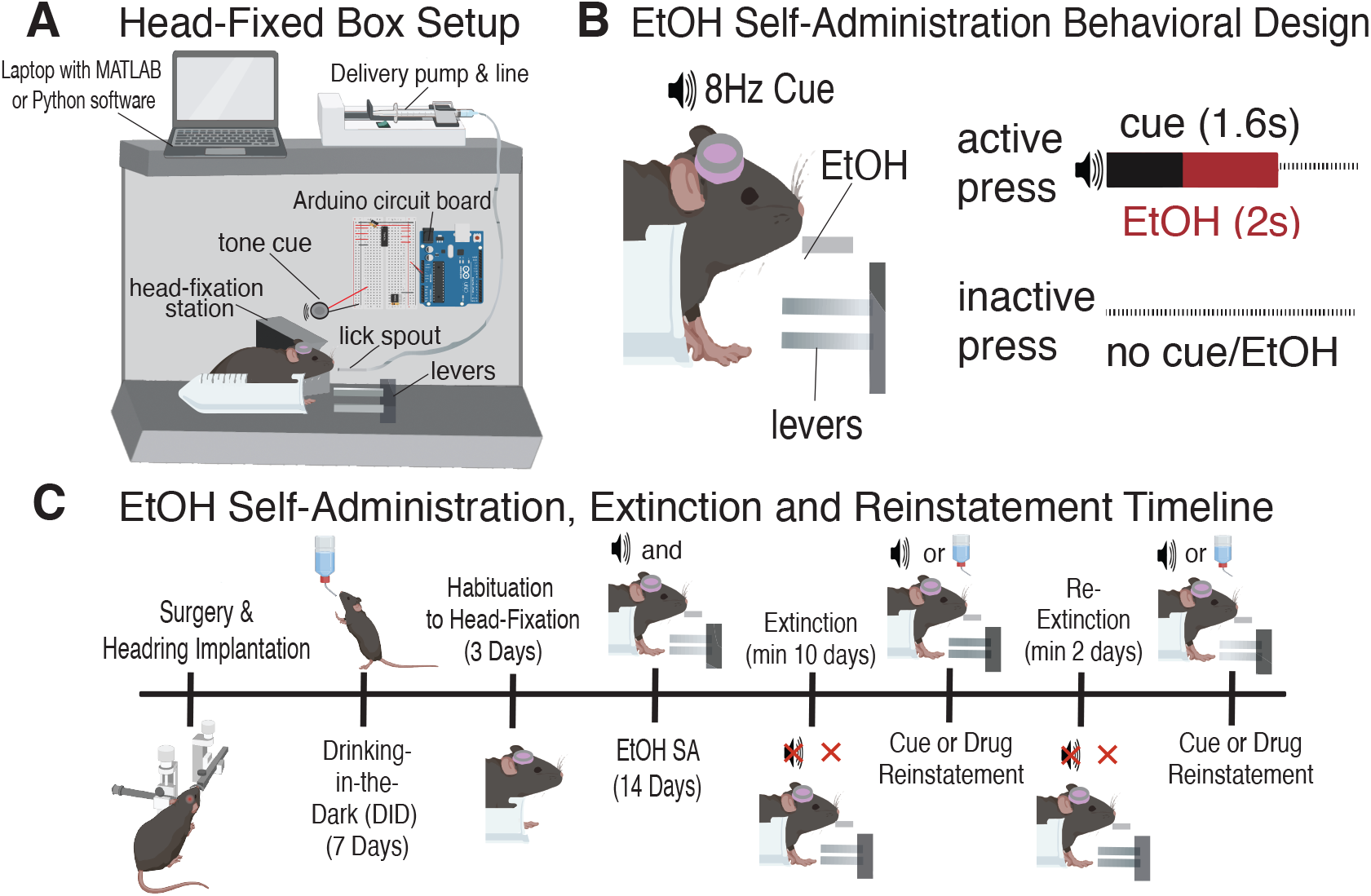
Setup and Timeline for Head-Fixed Self-Administration, Extinction and Reinstatement. (**A**) Diagram of head-fixed behavioral chambers (**B**) Schematic of behavioral design for acquisition of ethanol self-administration, wherein an active press results in a 1.6s cue presentation followed by a 15% ethanol reward and an inactive press result in no cue and no ethanol reward. (C) A timeline of head-fixed ethanol self-administration, extinction, and reinstatement procedures.

**Figure 2.**
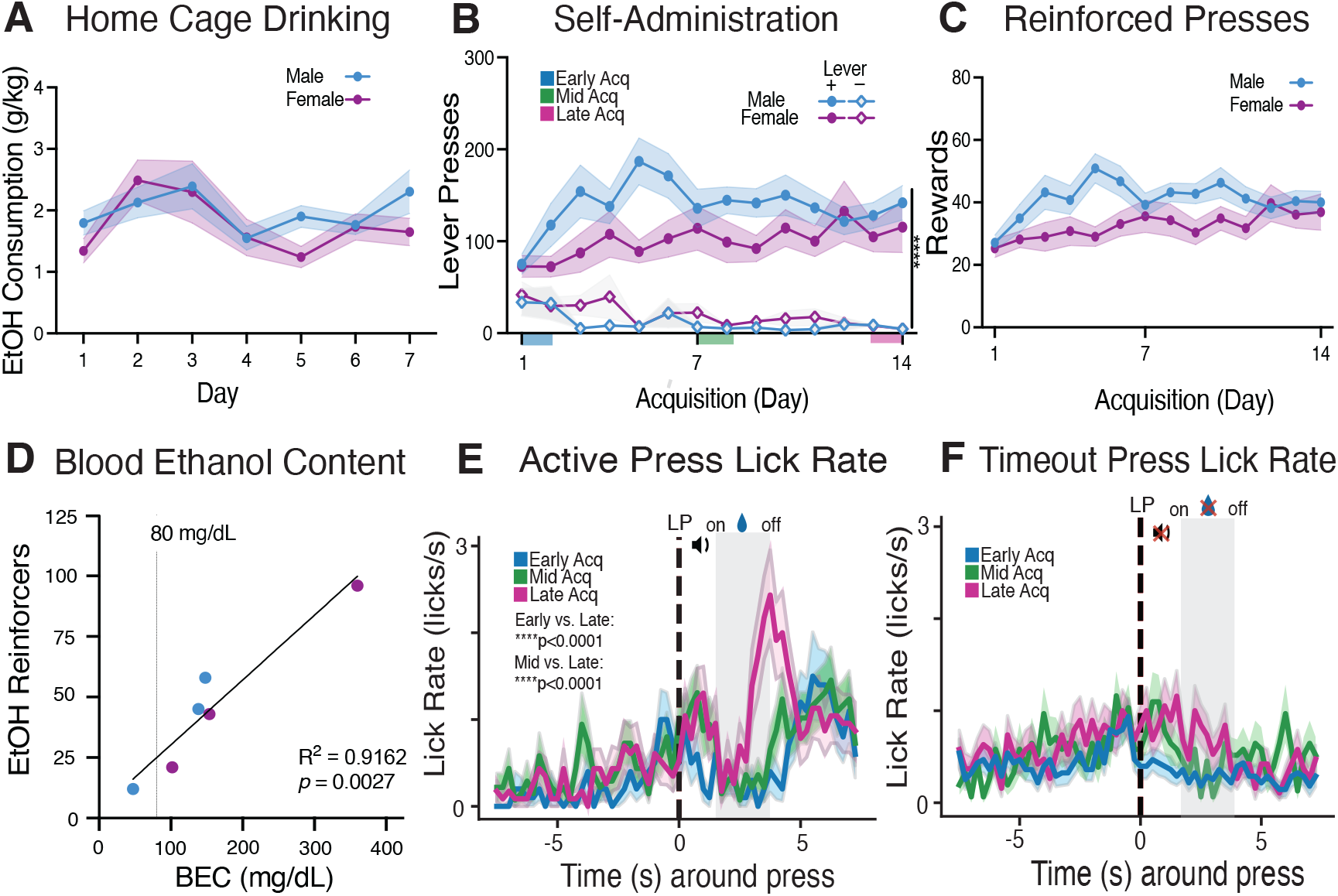
Head-Fixed Ethanol Self-Administration Paradigm. **A**) Home cage Drinking-in-the-Dark (DID) ethanol (15% v/v) consumption in females (purple) and males (blue). (**B**) Lever pressing behavior across 14 days of acquisition on the active and inactive levers for males and females showing significant discrimination between the active and inactive levers, \*\*\*\**p*<0.0001. (**C**) Reinforced active lever press responding across the course of behavioral acquisition. **(D)** Correlation of blood ethanol content to number of reinforcers during behavioral acquisition. (**E**) Moreover, when examining lick rate of animals during behavioral acquisition, we found that animals will have increased lick frequency to a reinforced active lever press across the course of learning, ****p<0.0001. (**F**) Furthermore, animals exhibit no changes in lick frequency in response to a timeout, non-reinforced lever press across the course of acquisition.

### Acquisition of Ethanol Self Administration

Following home cage DID exposure, mice were trained to press the active lever on an FR1 schedule of reinforcement for delivery of 15% (v/v) ethanol. Over the course of 14 days of acquisition animals learned to reliably discriminate between the active and inactive levers (**Fig 2B**). A mixed-effects three-way ANOVA showed a significant main effect of lever (but not day or sex, *p* = 0.6277 and 0.1344, respectively), such that overall mice pressed the active lever significantly more than the inactive lever, F_1,37_=120, *p*<0.0001. Additionally, there was no day x sex interaction (*p*=0.3661) nor a day x sex x lever interaction (*p*=0.2227), suggesting males and females performed comparably overall. However, there was a significant day x lever interaction, F_7.108,263.0_=3.779, *p*=0.0006, indicating significant discrimination between the active and inactive levers developed as acquisition went on, and a significant sex x lever interaction, F_1,37_=5.994, *p*=0.0192, suggesting males pressed the active lever slightly more than females. A two-way ANOVA analyzing number of ethanol reinforcers indicated a significant main effect of day F_13,481_=2.1, *p*=0.0092, suggesting that number of ethanol reinforcers earned increased across acquisition (**Fig 2C**). There was also a main effect of sex, F_1,37_=5.325, *p*=0.0267, suggesting that overall males earned more reinforcers than females (41.07 vs. 32.47, respectively. There was no significant day x sex interaction (*p*=0.1588). Following a subset of self-administration sessions, some animals received retro-orbital blood sampling to measure blood ethanol concentrations (BECs). The data evinced a strong positive correlation (r=0.957, p=0.0027) between the number of ethanol reinforcers earned and BECs achieved, revealing that animals become reliably intoxicated with an increasing number of ethanol reinforcers during acquisition (**Fig 2D**).

### Lick Rate During Acquisition of Ethanol Self-Administration

Lick rates were quantified in a subset of mice (n = 3-4 mice per day with at least 10 presses and 5 licks per trial) using a Python/Arduino user interface. To filter lick rates, the minimum lick duration was 1 ms and the maximum 70 ms; this filtering was designed to remove paw touches that may be longer than typical lick bouts. We conducted a two-way ANOVA analyzing lick rates across days (3 levels: Early, Mid, Late) and time (30 bins before and 30 bins after each press, with each bin representing 250 ms or total of 15 seconds analyzed). A two-way ANOVA (△lick rate) revealed a significant effect of day (F_(1,2)_ = 16.696, *p*<0.001), time (F_(1,59)_ = 6.754, *p*< 0.001) and a day x time interaction (F_(1,118)_ = 1.86, *p*<0.001) for licking behavior in response to a reinforced active lever press (**Fig 2E**). In contrast, mice did not increase licking rates during the equivalent ethanol delivery epoch when the active lever was pressed, but no ethanol or cue was presented (i.e., during timeout presses). Furthermore, a two-way ANOVA revealed a significant effect of day (F_(1,2)_ = 29.075, *p*<0.001) and time (F_(1,59)_ = 2.539, *p*< 0.001) (**Fig 2F**) for licking behavior in response to a timeout lever press. Moreover, post-hoc analyses were conducted using pairwise t-tests with Bonferroni correction. Post-hoc analyses examining infusion epoch (between 1.6-3.6s after lever press) showed significant differences in lick rate between early versus late acquisition (*p*<0.001) and mid versus late acquisition (*p*<0.001) in accordance to an active lever press. Further post-hoc analysis indicated no significant difference in lick rate between acquisition stages in response to a timeout lever press. Overall, data reveal that mice increase licking rates upon ethanol delivery across training.

### Extinction

After acquisition of ethanol self-administration, animals underwent extinction training (n=23; 13 males and 10 females). We have found that animals will sufficiently extinguish active lever pressing from the first day of extinction to the last day (**Fig 3A**). In the early days of extinction, animals will exhibit elevated lever pressing behavior that declines across time. A 2 × 2 RM ANOVA revealed a significant main effect of day, F_1,21_=9.120, *p*=0.0065, confirming that overall, animals significantly reduced lever-pressing behavior from the first day to the last day of extinction (**Fig 3A**). Additionally, a significant main effect of sex, F_1,21_=5.432, *p*=0.0298 and a significant day x sex interaction, F_1,21_=5.756, *p*=0.0258, revealed that males pressed the active lever on the first day of extinction significantly more than females (post-hoc *p*=0.0017), showing that males might be more prone to extinction burst behavior.

**Figure 3.**
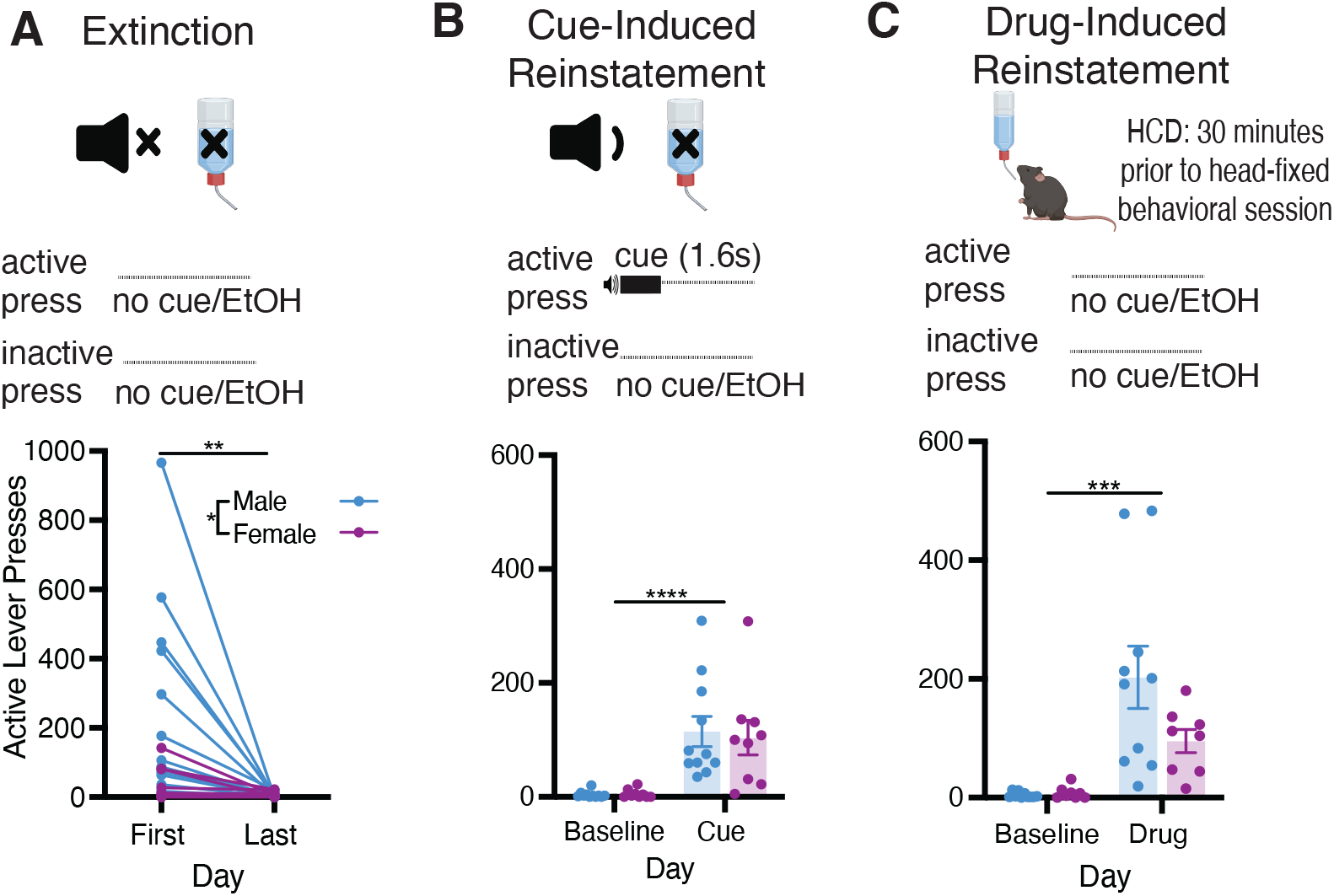
Extinction and reinstatement of operant responding for ethanol in a head-fixed paradigm. (**A**) During extinction learning, responses on the active lever no longer result in delivery of the tone cue or ethanol reward (top panel); males show significantly greater responding on day 1 of extinction compared to females (\**p*<0.05), but both male and female mice significantly reduce their lever pressing behavior from the first to last day of extinction (\*\**p*<0.01; bottom panel). (**B**) During cue-induced reinstatement testing, active lever results in delivery of the tone cue but no ethanol reward (top panel); both males and females show significantly greater responding on the cue-test day compared to their extinction baseline (\*\*\*\**p*<0.0001; bottom panel). (**C**) During drug-induced reinstatement testing, mice were given 30-min access to 15% ethanol in their home cage prior to placement in the head-fixed operant set up (top panel); both males and females show significantly greater responding on the drug-test day compared to their extinction baseline (\*\*\*\**p*<0.001; bottom panel).

### Reinstatement Tests Following Extinction

Lastly, we have found that head-fixed animals will reinstate ethanol-seeking behavior following extinction learning. Here we show that both male and female mice will equivalently reinstate active lever-pressing behavior to the tone cue previously paired with the ethanol reward (**Fig 3B**). Specifically, a 2 × 2 RM ANOVA revealed a significant main effect of day, F_1,18_=27.90, *p*<0.0001), such that active lever presses were increased after cue presentation in comparison to the previous day’s extinction baseline in a sex-independent manner (no main effect of sex, F_1,18_=0.0541, *p*=0.8187, and no day x sex interaction, F_1,18_=0.0974, *p*=0.7586). Moreover, these animals will reinstate lever pressing following 30 minutes of home cage drinking (HCD) prior to a head-fixed behavioral session. Using a 2 × 2 RM ANOVA we found that, similar to cue-induced reinstatement, both male and female mice will significantly reinstate active lever-pressing behavior in response to ethanol priming (**Fig 3C**) as shown by a main effect of day, F_1,16_=22.28, *p*=0.0002, but no main effect of sex, F_1,16_=2.777, *p*=0.1151, or a day x sex interaction, F_1,16_=3.377, *p*=0.0848.

## DISCUSSION

Collectively, our findings indicate that mice will self-administer ethanol in a head-fixed operant conditioning paradigm. Moreover, we find that animals will extinguish active lever-pressing in the absence of tone cue and ethanol reward. Importantly, when animals are re-introduced to a tone cue or ethanol reward, animals will reinstate active lever-pressing behavior. Therefore, we demonstrate that animals will readily acquire, extinguish and reinstate ethanol-seeking behaviors while head-fixed, effectively establishing the ability to apply novel methodologies such as simultaneous 2P imaging (Vollmer et al., 2021, Vollmer et al., 2022, Clarke et al., 2024, Kalelkar et al., 2024), providing for the first time a uniquely analogous behavioral paradigm to that in freely moving mice (Lopez and Becker, 2014, Meisch, 2001). These results indicate that our head-fixed ethanol self-administration paradigm provides a robust and novel technique to further dissect the mechanisms underlying ethanol seeking.

### Head-Fixation in Drug and Reward Self-Administration

Head-fixation during drug self-administration has been a reliable model to study various drugs of abuse, including heroin and cocaine (Vollmer et al., 2021, Paniccia et al., 2024, Clarke et al., 2024). Previous studies have found that animals will effectively acquire lever pressing, extinguish and reinstate during head-fixed heroin and cocaine self-administration (Vollmer et al., 2021). Notably, head-fixed animals will learn to discriminate between active and inactive levers, indicating the acquisition of goal-directed behavior that has been illustrated in freely-moving models. In addition to drug self-administration, the same operant self-administration, extinction, and reinstatement phenomena has been observed in sucrose reward-seeking models (Vollmer et al., 2021). Head-fixation during ethanol-seeking tasks is emerging as a novel strategy to examine circuitry during *in vivo* recordings (Timme et al., 2024, Kalelkar et al., 2024), providing a wealth of information on how ethanol can alter distinct neurons, circuits and ensembles in the brain during consumption. Other models have utilized aversion-resistance tasks during head-fixation (Timme et al., 2024) as well as other tasks such as operant licking tasks (Kalelkar et al., 2024) to examine how animals respond to various ethanol seeking contexts. However, the literature on head-fixed seeking tasks during ethanol consumption remains limited. Our data provide the first demonstration of a veritable operant-based ethanol-seeking task in a head-fixed preparation, which will enable the field to better investigate neural and circuit dysfunction during volitional ethanol-seeking and -taking behaviors.

### Integration of Head-Fixed Ethanol Self-Administration with 2-Photon Microscopy

Head-fixation provides stability that facilitates 2P imaging in awake animals, enabling high-resolution investigation of neuronal activity during behavior. Two-photon microscopy has become a critical tool for studying neuropsychiatric disorders, including substance use disorders, by allowing deep tissue imaging with single-cell specificity (Grienberger et al., 2022, Svoboda and Yasuda, 2006). Two-photon microscopy utilizes two near-infrared photons to excite fluorophores within a sample simultaneously, allowing for fluorescence detection despite the longer excitation photo wavelength (Svoboda and Yasuda, 2006). This excitation facilitates tissue penetration and minimizes background noise, markedly enhancing imaging resolution, clarity, and potential depth as compared with other *in vivo* imaging applications. Moreover, 2P microscopy reduces phototoxicity and photobleaching, which enables prolonged cellular and molecular observations *in vivo* (Grienberger et al., 2022).

A key advantage of 2P microscopy is it enables tracking of individual cellular elements over extended periods of time (weeks to months; (Namboodiri et al., 2019, Rossi et al., 2019) providing unique insight into neuroplasticity associated with various neurobiological conditions. This temporal tracking is highly multifaceted in that it may be used to track single cells (Rossi et al., 2019); dendrites (Sadakane et al., 2015), dendritic spines (Pan and Gan, 2008, Yu and Zuo, 2014), and axons (Davalos et al., 2008). Such longitudinal perspective will be particularly valuable for studying AUD, where dynamic changes in neuronal activity and connectivity underlie the progression from initial ethanol exposure to compulsive seeking (Timme et al., 2022, Radke et al., 2017). Combining head-fixed ethanol self-administration with two-photon microscopy allows the application of single-cell optogenetics (Piantadosi et al., 2024, Yang and Yuste, 2018), enabling functional targeting of neuronal clusters and populations that may be implicated in ethanol seeking behaviors. Furthermore, spatial sequencing enables the integration of two-photon imaging datasets with RNA sequencing, providing unprecedented insight into the computational dynamics of transcriptomically unique cell populations (Xu et al., 2020). While some have demonstrated the benefits of using head-fixed models in conjunction with two-photon imaging and ethanol-related tasks (Kalelkar et al., 2024), research combining 2P microscopy with ethanol-seeking tasks remains limited. By offering a dynamic, *in vivo* perspective concerning the neural adaptations associated with the development of AUD, this technique could vastly improve the dissection of the complex neuronal circuits that longitudinally adapt to functionally guide dysfunctional alcohol use and seeking behaviors.

## FUTURE DIRECTIONS AND LIMITATIONS

While our head-fixed model has shown robust and consistent results in ethanol self-administration, there are several important considerations for adapting freely-moving behaviors to head-fixed models, though our data suggest these concerns can be largely overcome. One of the primary concerns with head-fixed behavioral tasks is potential stress responses that may result from head restraint (Juczewski et al., 2020). This presents a valid concern as stress responses affect a variety of behaviors, including drug seeking (Mantsch and Katz, 2007). However, longevity of habituation to the head-fixed setup may allow rodents to become more acclimated to this restraint and prevent stress levels from influencing behavior (Schwarz et al., 2010). Moreover, previous studies implementing head-fixed drug self-administration have not indicated changes in a behavior as a result of head-restraint (Vollmer et al., 2021) and remain sensitive to stress (Vollmer et al., 2022). Nevertheless, as technology and equipment improve, there has been increasing integration of elements such as a running wheel or a treadmill into head-fixed setups to track movement and limit potential stress from restraint. Furthermore, additions to our current head-fixed setup could be added to track other metrics during operant tasks, such as a camera to measure aspects such as pupil diameter and whisker movement, which could be used to assess states of arousal (Ganea et al., 2020) and object recognition (Adibi, 2019).

## CONCLUSION

Collectively, our results show that head-fixed animals will rapidly and reliably learn goal-directed behaviors to seek ethanol. Our head-fixed paradigm indicates that animals will learn to discriminate between active and inactive levers, effectively extinguish ethanol seeking behavior and subsequently reinstate to ethanol and ethanol-paired cues. These results indicate that self-administration protocols can be effectively applied to head-fixed mice during 2P imaging, revealing unparalleled access to the brain during ethanol seeking behaviors.

## AUTHOR CONTRIBUTIONS

ALW, JMO and JAR designed the experiments and wrote the manuscript. ALW, KTW, SB, CJC, REC, MSB, and MRM helped conduct the experiments. SB, REC, MSB, MRM, KMV, JEP, EMD, RIG, JB, JB, LMM, AMW, LMG and BEP provided intellectual support and training for self-administration studies. All authors contributed to the article and approved the final submitted version.

## ACKNOWLEDGEMENTS

We would like to thank Drs. Howard C. Becker, Jacqueline F. McGinty and John J. Woodward for intellectual support and advice.

